# Enhanced Killing of Triple-Negative Breast Cancer Cells by Reassortant Reovirus and Topoisomerase Inhibitors

**DOI:** 10.1101/644815

**Authors:** Roxana M. Rodríguez Stewart, Jameson T.L. Berry, Angela K. Berger, Sung Bo Yoon, Jaime A. Guberman, Nirav B. Patel, Gregory K. Tharp, Steven E. Bosinger, Bernardo A. Mainou

**Affiliations:** Emory University School of Medicine, Atlanta, GA 30322; Department of Pediatrics, Emory University School of Medicine, Atlanta, GA 30322; Children’s Healthcare of Atlanta, Atlanta, GA, 30322; Emory Vaccine Center, Yerkes National Primate Research Center, Atlanta, GA 30322

**Author notes:** R.M.R.S. and J.T.L.B. contributed equally to this work. To whom correspondence should be addressed: Emory University School of Medicine, Department of Pediatrics, ECC 564, 2015 Uppergate Drive, Atlanta, GA 30322.

## Abstract

Breast cancer is the second-leading cause of cancer-related deaths in women in the United States. Triple-negative breast cancer constitutes a subset of breast cancer that is associated with higher rates of relapse, decreased survival, and limited therapeutic options for patients afflicted with this type of breast cancer. Mammalian orthoreovirus (reovirus) selectively infects and kills transformed cells and a serotype 3 reovirus is in clinical trials to assess its efficacy as an oncolytic agent against several cancers. It is unclear if reovirus serotypes differentially infect and kill triple-negative breast cancer cells and if reovirus-induced cytotoxicity of breast cancer cells can be enhanced by modulating the activity of host molecules and pathways. Here, we generated reassortant reoviruses by forward genetics with enhanced infective and cytotoxic properties in triple-negative breast cancer cells. From a high-throughput screen of small molecule inhibitors, we identified topoisomerase inhibitors as a class of drugs that enhance reovirus infectivity and cytotoxicity of triple-negative breast cancer cells. Treatment of triple-negative breast cancer cells with topoisomerase inhibitors activates DNA damage response pathways and reovirus infection induces robust production of Type III, but not Type I, interferon. Together, these data show that reassortant viruses with a novel genetic composition generated by forward genetics in combination with topoisomerase inhibitors more efficiently infect and kill triple-negative breast cancer cells.

## Importance

Patients afflicted by triple-negative breast cancer have decreased survival and limited therapeutic options. Reovirus infection results in cell death of a variety of cancers, but it is unknown if different reovirus types lead to triple-negative breast cancer cell death. In this study, we generated two novel reoviruses that more efficiently infect and kill triple-negative breast cancer cells. We show that infection in the presence of DNA-damaging agents enhances infection and triple-negative breast cancer cell killing by reovirus. These data suggest that a combination of a genetically engineered oncolytic reovirus and topoisomerase inhibitors may provide a potent therapeutic option for patients afflicted with triple-negative breast cancer.

## Introduction

Breast cancer is the leading cause of cancer and second leading cause of deaths by cancer in women in the United States (1). Triple-negative breast cancer (TNBC) constitutes approximately 15% of breast cancers, has a higher rate of relapse, and shorter overall survival after metastasis than other subtypes of breast cancer (2). TNBC more frequently affects the young, is more prevalent in African American women, and tumors are larger in size and biologically more aggressive (3). TNBC is characterized by the lack of expression of estrogen receptor (ER), progesterone receptor (PR), and human epidermal growth factor receptor 2 (HER2/neu) and can be classified into seven subtypes based on their genetic signature (3). Although targeted therapies against hormone receptor-positive and HER2-positive breast cancer have been efficacious, the absence of these molecules on TNBC cells has limited treatment to cytotoxic chemotherapy, radiotherapy, and surgery (4, 5). This raises a need for targeted therapeutics against this type of cancer.

The concept that viruses can promote tumor regression is nearly as old as the discovery of viruses (6). The deregulated expression of viral receptors, endocytic uptake molecules, proteases, altered metabolic states, and impaired innate immunity make cancer cells ideally suitable for virus infection and replication (7–9). In addition to directly impacting cancer cell biology, oncolytic viruses can elicit anti-tumor immune responses and serve as adjuvants for other cancer therapies (10–12). Several viruses are under study to assess their oncolytic properties against several cancers (7, 8). Nonfusogenic mammalian orthoreovirus (reovirus, Reolysin) is a non-enveloped double-stranded RNA (dsRNA) virus in the *Reoviridae* family. A serotype 3 reovirus is in Phase I and II clinical trials (clinicaltrials.gov: NCT01622543, NCT01656538) to assess its efficacy against a variety of cancers (13). Reovirus can be delivered to patients via intratumoral and intravenous administration and can be effective in combination therapy (14). Reovirus has an inherent preference to replicate in tumor cells, making it ideally suited for use in oncolytic virotherapies (15, 16). However, the cellular and viral factors that promote preferential reovirus infection of cancer cells are not fully elucidated.

Reovirus has a segmented genome with three large (L), three medium (M), and four small (S) dsRNA gene segments (17). There are three different reovirus serotypes (Type 1, 2, and 3) based on the neutralization ability of antibodies raised against the σ1 attachment protein that is encoded by the S1 gene segment (18, 19). Reoviruses infect most mammals, and although humans are infected during childhood, infection seldom results in disease (18, 20-22). Reovirus induces programmed cell death *in vitro* and *in vivo* (23–30). Although both Type 1 and Type 3 reovirus can induce apoptosis, Type 3 reoviruses induce apoptosis and necroptosis more efficiently in most cells (18, 23, 24). Serotype-dependent differences in apoptosis induction segregate with the S1 and M2 gene segments (31–33). However, there is a limited understanding of the viral factors that determine preferential replication and killing of cancer cells.

In this study, we show that co-infection and serial passaging of parental reoviruses in TNBC cells yields reassortant viruses with enhanced oncolytic capacities compared to parental reoviruses. Reassortant reoviruses have a predominant Type 1 genetic composition with some Type 3 gene segments as well as synonymous and non-synonymous point mutations. We show that reassortant reoviruses have enhanced infective and cytotoxic capacities in TNBC cells compared to parental viruses. To further enhance the oncolytic properties of these reassortant viruses, we used a high-throughput screen of small molecule inhibitors and identified DNA-damaging topoisomerase inhibitors as a class of drugs that reduces TNBC cell viability while enhancing reovirus infectivity. Infection of TNBC cells in the presence of topoisomerase inhibitors results in induction of DNA damage, increased levels of Type III but not Type I interferon, and enhanced cell killing. Together, we show that reassortant reoviruses with a novel genetic composition have enhanced oncolytic properties and pairing of topoisomerase inhibitors with reovirus potentiates TNBC cell killing.

## Results

### Generation of reassortant viruses in triple-negative breast cancer cells by forward genetics

Reovirus serotypes have distinct infective, replicative, and cell killing properties and the segmented nature of the reovirus genome allows the generation of viruses with novel properties through gene reassortment following co-infection (34, 35). To generate reoviruses with enhanced replicative properties in TNBC cells, MDA-MB-231 cells were co-infected with prototype laboratory strains T1L, T2J, and T3D and serially passaged in these cells ten or twenty times (FIG 1A). Following serial passage, individual viral clones were isolated by plaque assay and the gene segment identity for each clone (44 clones following 10 passages, 45 clones following 20 passages) was determined by SDS-gel electrophoresis (FIG 1B). Of the 44 isolates analyzed following 10 serial passages, 8 distinct electropherotypes were identified, with 23 isolates (52%) having the same electropherotype (r2Reovirus) (FIG 1C). Following 20 serial passages, 6 distinct isolates were identified, including two (r9 and r10) that were not observed after passage 10 (FIG 1D). The most predominant electropherotypes following 20 serial passages were r1Reovirus and r2Reovirus, constituting 33% and 27% respectively of all isolates. Illumina Next-Generation Sequencing (NGS) revealed that r1Reovirus is composed of seven gene segments from T1L and three from T3D (L2, M2, S2), while r2Reovirus is composed of nine gene segments from T1L and one from T3D (M2) (FIG 2). In addition, both viruses have nonsynonymous point mutations that result in an Ala to Thr substitution at amino acid 160 in L3, an Ile to Val substitution at amino acid 250 in S3, and Val to Ile substitution at amino acid 49 in S4. A Val to Ile substitution at amino acid 426 in S2 and a Pro to Thr substitution at amino acid 160 are also found in r1Reovirus. In addition, r1Reovirus and r2Reovirus have several synonymous point mutations (Table S1). The r1Reovirus S2 gene segment, but no other gene segment, has single residue variations that range from 35% to 65%. This suggests that r1Reovirus may not be clonal but a mixture of two or more viruses with different genetic signatures at the S2 gene segment or that the virus contains two S2 gene segments. We did not detect single residue variations in gene segments from either parental T1L, T2J, or T3D or r2Reovirus, suggesting this is not an intrinsic property of the S2 gene segment carried from parental viruses. Together, these data indicate that co-infection and serial passaging of reoviruses in MDA-MB-231 cells leads to the generation of reassortant reoviruses with novel genetic compositions.

**FIG 1.**
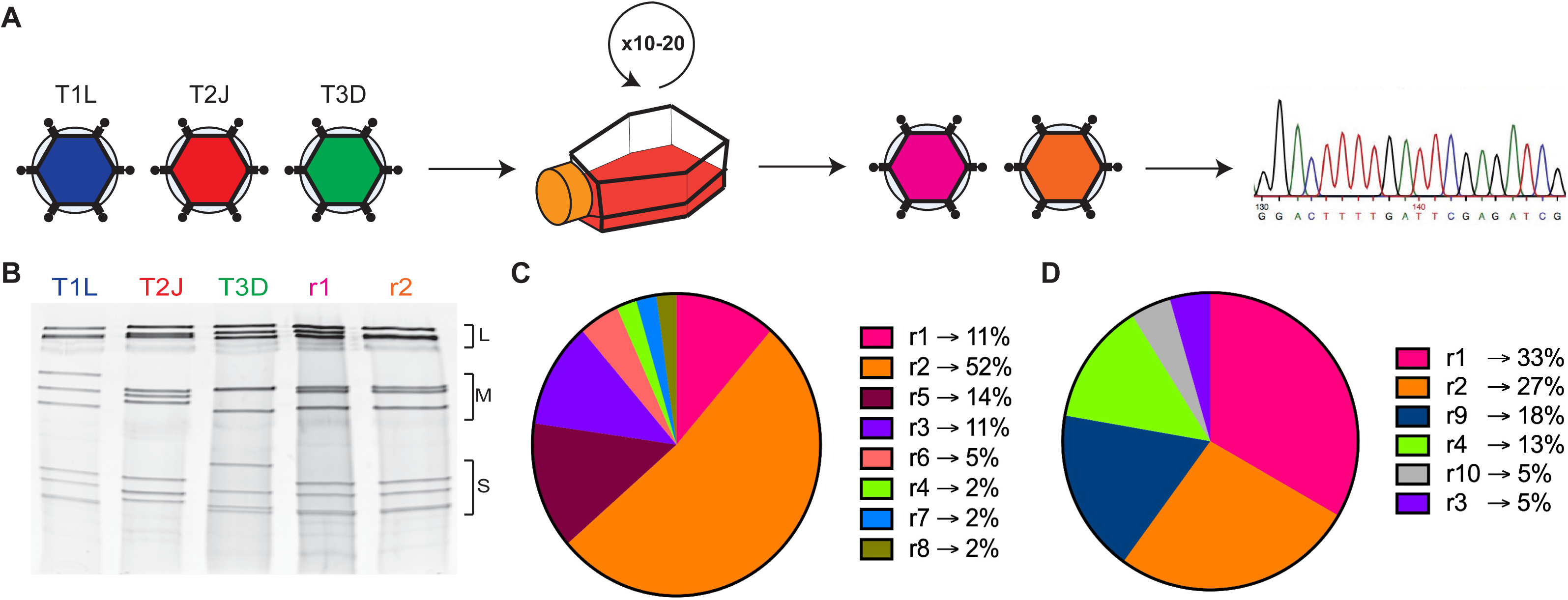
Generation of reoviruses by forward genetics in MDA-MB-231 cells. (A) Triple-negative breast cancer MDA-MB-231 cells were co-infected with T1L, T2J, and T3D and serially passaged ten or twenty times. Virus isolates were obtained following plaque assay on L929 cells and sequenced by Illumina Next-Generation Sequencing. (B) Polyacrylamide gel electrophoresis of reovirus parental strains T1L, T2J, and T3D and r1Reovirus (r1) and r2Reovirus (r2). Strains are differentiated by migration patterns of three large (L), three medium (M), and four small (S) gene segments. (C) Percentage of viral isolates with a specific electropherotype following 10 serial passages in MDA-MB-231 cells (n = 44). r1Reovirus (pink) accounts for 11% of isolates while r2Reovirus (orange) accounts for 52%. (D) Percentage of viral isolates with a specific electropherotype following 20 serial passages in MDA-MB-231 cells (n = 45). r1Reovirus (pink) accounts for 33% of isolates while r2Reovirus (orange) accounts for 27%.

**FIG 2.**
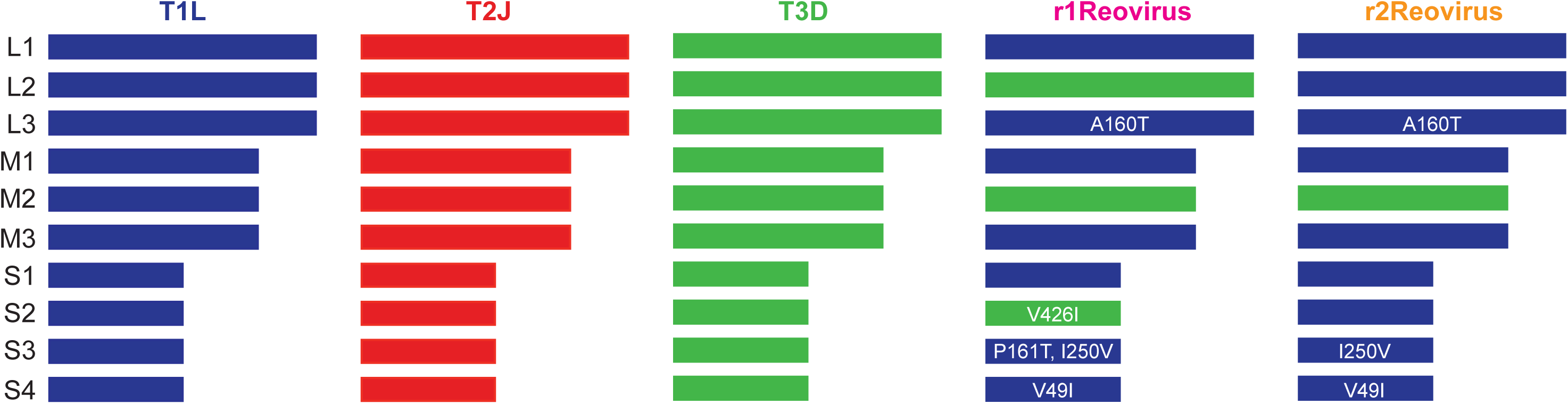
Genetic composition of r1Reovirus and r2Reovirus. The genetic composition of parental and reassortant r1Reovirus and r2Reovirus was determined by Illumina Next-Generation sequencing. r1Reovirus has seven gene segments from T1L and three from T3D (S2, M2, L2) and five nonsynonymous point mutations (L3 A160T, S2 V426I, S3 P161T, I250V, and S4 V49I). r2Reovirus has nine gene segments from T1L and one from T3D (M2) and three nonsynonymous point mutations (L3 A160T, S3 I250V, and S4 V49I). Both r1Reovirus and r2Reovirus have several synonymous point mutations.

### Reassortant reoviruses attach to cells with similar efficiency but infect MDA-MB-231cells more efficiently than parental reoviruses

Reovirus attaches to cells via a strength-adhesion mechanism whereby the viral attachment fiber σ1 binds to cell-surface carbohydrate and proteinaceous receptors JAM-A or NgR1 (36–40). To determine the attachment efficiency of r1Reovirus and r2Reovirus in comparison to parental reoviruses, MDA-MB-231 cells were adsorbed with vehicle (mock) or Alexa 633 (A633)-labeled T1L, T3D, T3C$ (the reovirus strain currently in clinical trials), or reassortant reoviruses at an MOI of 5×10^4^ particles/cell and assessed for cell surface reovirus by flow cytometry (FIG 3A). Reassortant reoviruses attach to cells with similar efficiency as T1L, but less efficiently than Type 3 reoviruses T3D and T3C$. As reassortant reoviruses contain a T1L S1 gene segment, it is not surprising that they attach to cells to similar levels as parental T1L. These data also indicate that other genetic changes found in r1Reovirus and r2Reovirus do not impact the ability of the viruses to attach to cells.

**FIG 3.**
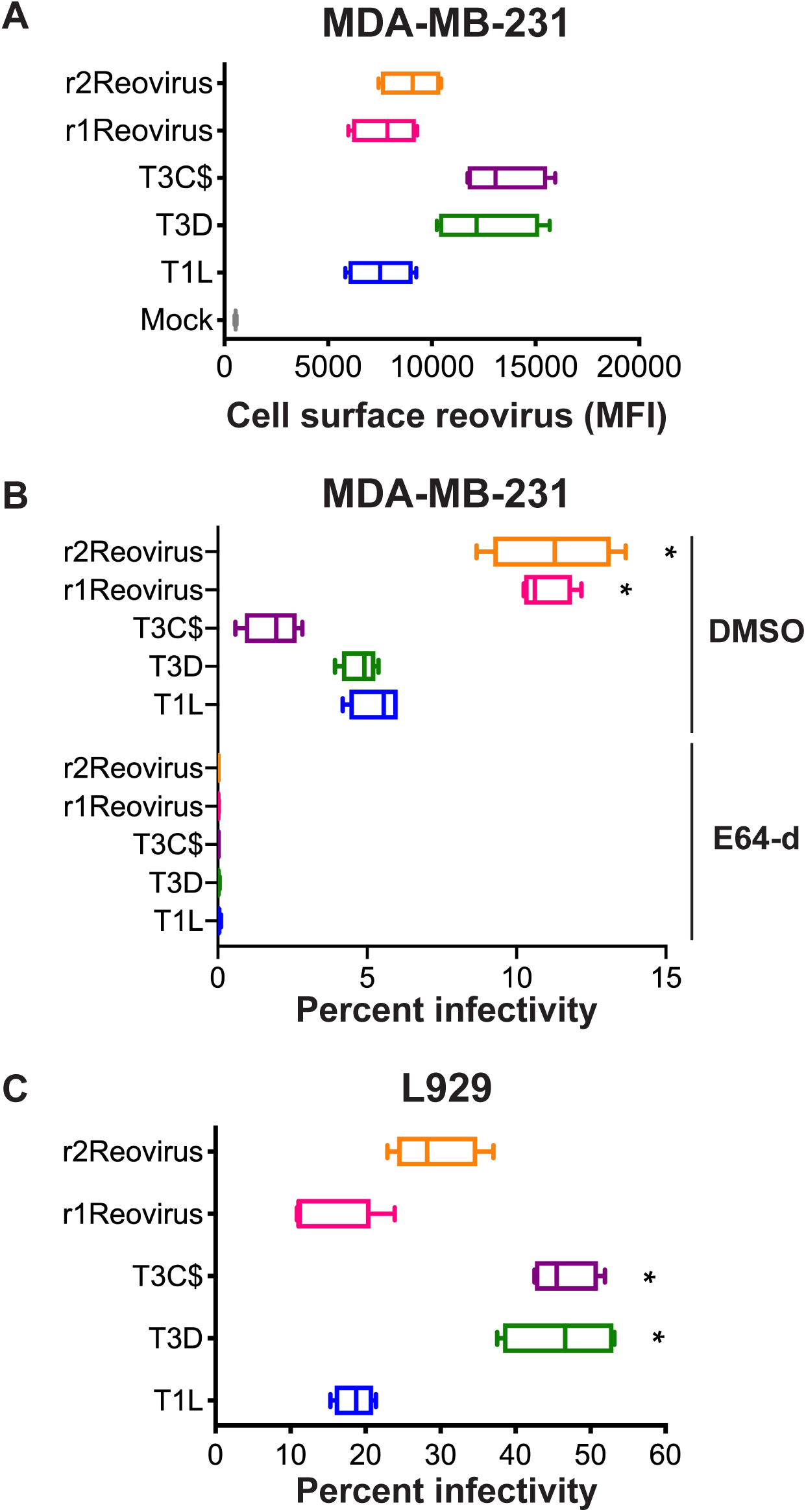
Attachment and infectivity of MDA-MB-231 cells by reassortant reoviruses. (A) MDA-MB-231 cells were adsorbed with A633-labeled T1L, T3D, T3C$, or reassortant reoviruses at an MOI of 5x10^4^ particles/cell and assessed for cell-surface reovirus by flow cytometry. Results are expressed as box and whisker plots of cell surface reovirus mean fluorescence intensity (MFI) for quadruplicate independent experiments. (B) MDA-MB-231 cells were treated with DMSO or 4 μM E64-d and adsorbed with T1L, T3D, T3C$, or reassortant reoviruses at an MOI of 100 PFU/cell and assessed for infectivity after 18 h by indirect immunofluorescence using reovirus-specific antiserum. (C) L929 cells were adsorbed with T1L, T3D, T3C$, or reassortant reoviruses at an MOI of 5 PFU/cell and assessed for infectivity after 18 h by indirect immunofluorescence using reovirus-specific antiserum. Results are expressed as box and whisker plots of percent infectivity for quadruplicate independent experiments. *, *P* ≤ 0.0003, **, *P* < 0.0001 in comparison to T1L by two-way ANOVA with Tukey’s multiple comparison test.

To determine how genetic changes in r1Reovirus and r2Reovirus affect reovirus infection of TNBC cells, MDA-MB-231 cells were pretreated with DMSO or the cysteine protease inhibitor E64-d, which blocks reovirus cell entry by preventing proteolysis during endocytic uptake (41), adsorbed with mock, T1L, T3D, T3C$, or reassortant reoviruses at an MOI of 100 PFU/cell and assessed for infectivity after 18 h by indirect immunofluorescence using reovirus-specific antiserum (FIG 3B). In contrast to that observed in the attachment assay, r1Reovirus and r2Reovirus infect MDA-MB-231 cells more efficiently than parental reoviruses or T3C$, with both reassortant viruses infecting cells over 2-fold more efficiently. Infection with all viruses tested was impaired by E64-d, indicating a similar requirement for proteolytic processing during entry. These data indicate that reassortant reoviruses establish infection more efficiently in MDA-MB-231 cells than parental reoviruses and that infection of these cells requires proteasomal processing of the virion during cell entry.

To determine if the increased infectivity of the reassortant viruses is limited to MDA-MB-231 cells, the infectivity of parental and reassortant reoviruses was assessed on murine L929 fibroblasts, which are highly susceptible to reovirus infection and are used to propagate the virus (FIG 3C). L929 cells were adsorbed with mock, T1L, T3D, T3C$, or reassortant reoviruses at an MOI of 5 PFU/cell and assessed for infectivity after 18 h by indirect immunofluorescence using reovirus-specific antiserum. In contrast to that observed in MDA-MB-231 cells, reassortant reoviruses infect L929 cells to similar levels as parental T1L, but less efficiently than both T3D and T3C$. These data indicate that r1Reovirus and r2Reovirus more efficiently infect TNBC cells, but not L929 cells. This suggests that the genetic changes found in the reassortant viruses confer enhanced infection in the TNBC cells used for serial passage at a step after attachment.

### Replication kinetics of reassortant reoviruses are similar to T1L but faster than T3D

To determine the replication efficiency of parental and reassortant reoviruses, MDA-MB-231 cells were adsorbed with mock, T1L, T3D, T3C$ or reassortant reoviruses at an MOI of 10 PFU/cell and assessed for viral replication over a 3 day course of infection (FIG 4). Despite the differences observed in infectivity, all viruses except T3D replicated with similar kinetics, with T3C$ having slightly faster replication kinetics at day 1 post infection than all other viruses tested (FIG 4B). T1L, T3C$, r1Reovirus, and r2Reovirus had similar replication kinetics at days 2 and 3 post infection. T3D replication kinetics were significantly slower than all other viruses tested. Interestingly, although T3C$ only differs from T3D by 22 amino acids, its replication kinetics are more similar to T1L and the reassortant reoviruses than T3D. These data indicate that although reassortant reoviruses establish infection in MDA-MB-231 cells more efficiently than parental reoviruses, replication kinetics are similar to T1L but significantly enhanced compared to T3D.

**FIG 4.**
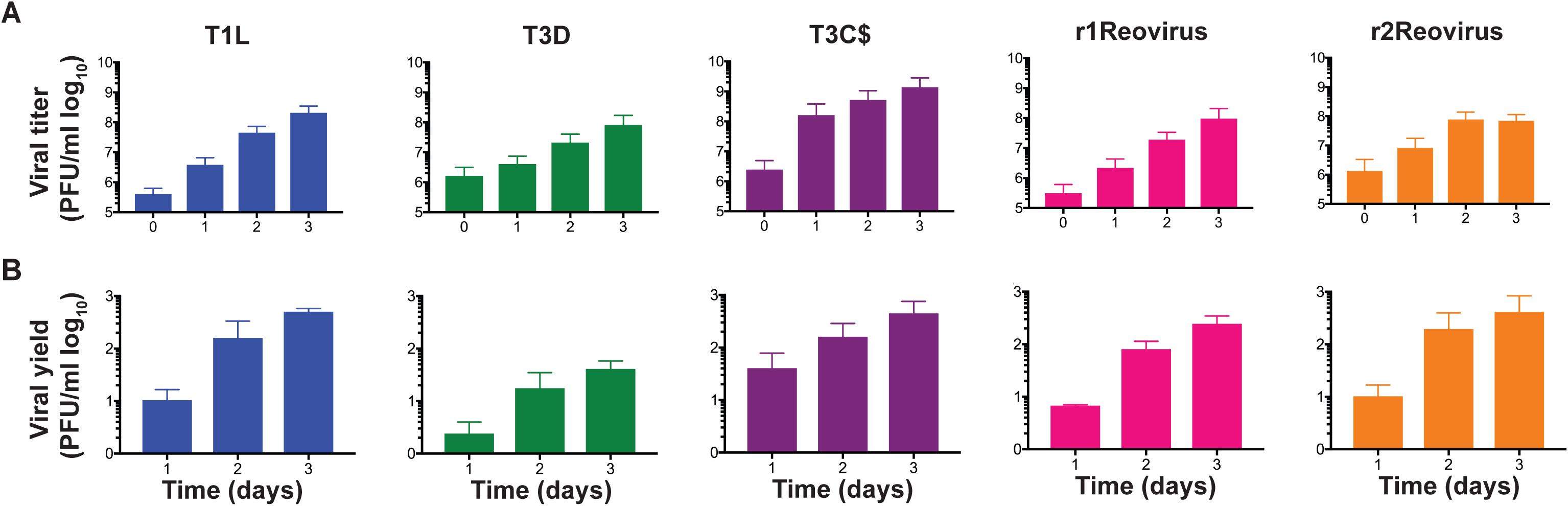
Reassortant viruses replicate with similar kinetics than T1L and T3C$, but faster than T3D, in MDA-MB-231 cells. T1L, T3D, T3C$, r1Reovirus, and r2Reovirus were adsorbed at an MOI of 10 PFU/cell and (A) viral titers and (B) viral yields were determined by plaque assay on L929 cells at 0-3 days post infection. The results are presented as (A) mean viral titers (± SEM) or (B) mean viral yields (± SEM) from day 0 post infection.

### r1Reovirus and r2Reovirus impact cell viability with faster kinetics than parental reoviruses in MDA-MB-231 but not L929 cells

Type 3 reoviruses induce cell death more efficiently than Type 1 reoviruses *in vitro* and *in vivo* and T3C$ is the reovirus strain currently in clinical trials to test its efficacy as an oncolytic against a variety of cancers (33, 42). To determine the efficacy of viral-induced cytotoxicity in TNBC cells, MDA-MB-231 cells were adsorbed with mock, T1L, T3D, T3C$, r1Reovirus and r2Reovirus at an MOI of 500 PFU/cell, or treated with staurosporine as a positive control, and assessed for cell viability for 7 days (FIG 5A). Compared to mock-infected cells, all reoviruses tested impaired cell viability, with reassortant reoviruses impairing cell viability with the fastest kinetics. In reassortant reovirus-infected cells, cell viability peaked at day 2 post infection, reaching levels similar to staurosporine by day 5 post infection. Cell viability peaked at day 3 post infection in T1L-, T3D-, and T3C$-infected cells reaching staurosporine levels by day 5 with T1L and day 6 with T3C$. T3D-infected cells did not reach staurosporine levels during the time course. Overall, the impact on cell viability by reassortant viruses was 1 day ahead of T1L and T3C$ and 2-3 days ahead of T3D. To determine if similar effects on cell viability could be observed in another TNBC cell line, MDA-MB-436 cells were infected with mock, T1L, T3D, T3C$, or r2Reovirus and assessed for cell viability over 6 days (FIG 5B). Similar to that observed in MDA-MB-231 cells, r2Reovirus induced cell death with faster kinetics than either parental T1L or T3D, or T3C$. T3D did not significantly impact MDA-MB-436 cell viability.These data show that reassortant viruses negatively affect cell viability of TNBC cells more efficiently than parental reoviruses and the oncolytic T3C$ strain. These data also suggest that T3D is not efficient at inducing cell death in at least a subset of TNBC cells.

**FIG 5.**
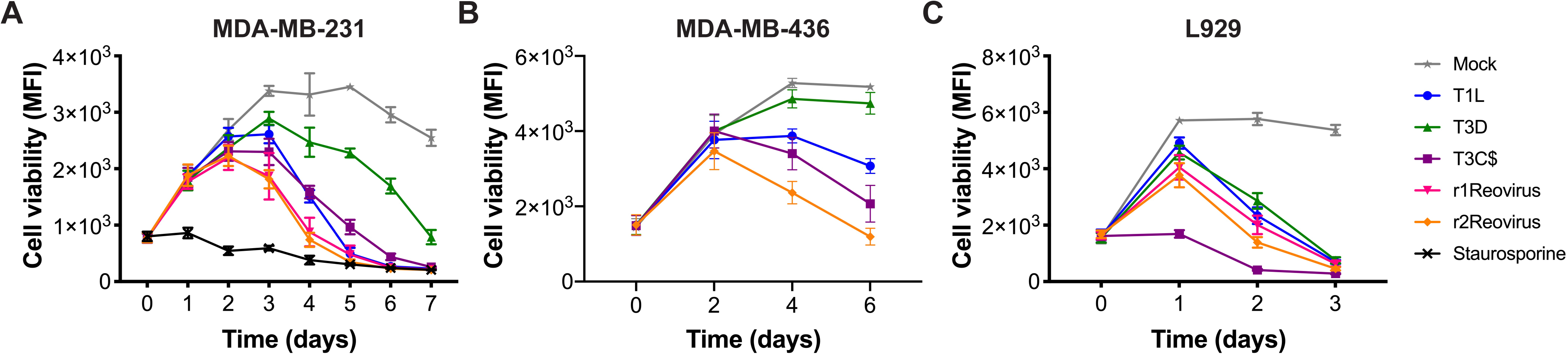
Impact on cell viability of TNBC cells and L929 cells following reovirus infection. (A) MDA-MB-231, (B) MDA-MB-436, and (C) L929 cells were adsorbed with T1L, T3D, T3C$, r1Reovirus, or r2Reovirus at an MOI of 500 PFU/ml or treated with 1 μM staurosporine and cell viability was assessed at times shown. Results are presented as mean fluorescence intensity (MFI) and SEM for four independent experiments.

To determine if r1Reovirus and r2Reovirus differ from parental reoviruses in their ability to impair cell viability of non-TNBC cells, L929 cells were adsorbed with mock, T1L, T3D, T3C$, r1Reovirus, or r2Reovirus at an MOI of 500 PFU/cell and assessed for cell viability over a 3 day time course (FIG 5C). In contrast to that observed in MDA-MB-231 cells, all reoviruses tested impaired cell viability with relatively similar kinetics except for T3C$, which impaired L929 cell viability with significantly faster kinetics. These data indicate that reassortant viruses induce cell death with faster kinetics in MDA-MB-231 and MDA-MB-436 cells but not in L929 cells. As r2Reovirus had enhanced infectivity and cytotoxicity in MDA-MB-231 compared to parental viruses and no single nucleotide variants in the S2 gene segment, experiments in the rest of the study were performed with r2Reovirus.

### Identification of small molecules that impact reovirus infectivity of MDA-MB-231 cells

To identify small molecule inhibitors that enhance the oncolytic potential of reovirus, a high-throughput screen to assess the effect of small molecules from the NIH Clinical Collection I and II (NCC) on reovirus infectivity was performed. The NCC is composed of compounds that have been through Phase I-III clinical trials. To test the effects on reovirus infectivity of compounds in the NCC, MDA-MB-231 cells were pre-treated with vehicle (DMSO), 4 μM E64-d, or 10 μM NCC compounds for 1 h. r2Reovirus was added to cells at an MOI of 20 PFU/cell, incubated for 20 h post infection in the presence of DMSO, 2 μM E64-d, or 5 μM NCC compounds, and scored for infectivity by indirect immunofluorescence using reovirus-specific antiserum (FIG 6A, Table S2). Of the 700 compounds in the NCC, 20 increased reovirus infectivity whereas 17 decreased infectivity (FIG 6B). Six microtubule-inhibiting compounds impaired reovirus infectivity, corroborating a need for microtubule function in reovirus cell entry (43). The sodium ATPase pump inhibitor digoxin and two serotonin antagonists also impaired reovirus infection, corroborating a role for the sodium ATPase pump and serotonin receptors in reovirus infection (44, 45). Four topoisomerase inhibitors, doxorubicin, epirubicin, etoposide (topoisomerase II inhibitors) and topotecan (topoisomerase I inhibitor), significantly enhanced reovirus infectivity. Topoisomerase inhibitors can sensitize TNBC cells to cell death but it is unknown how they impact reovirus-mediated cell death (46).

**FIG 6.**
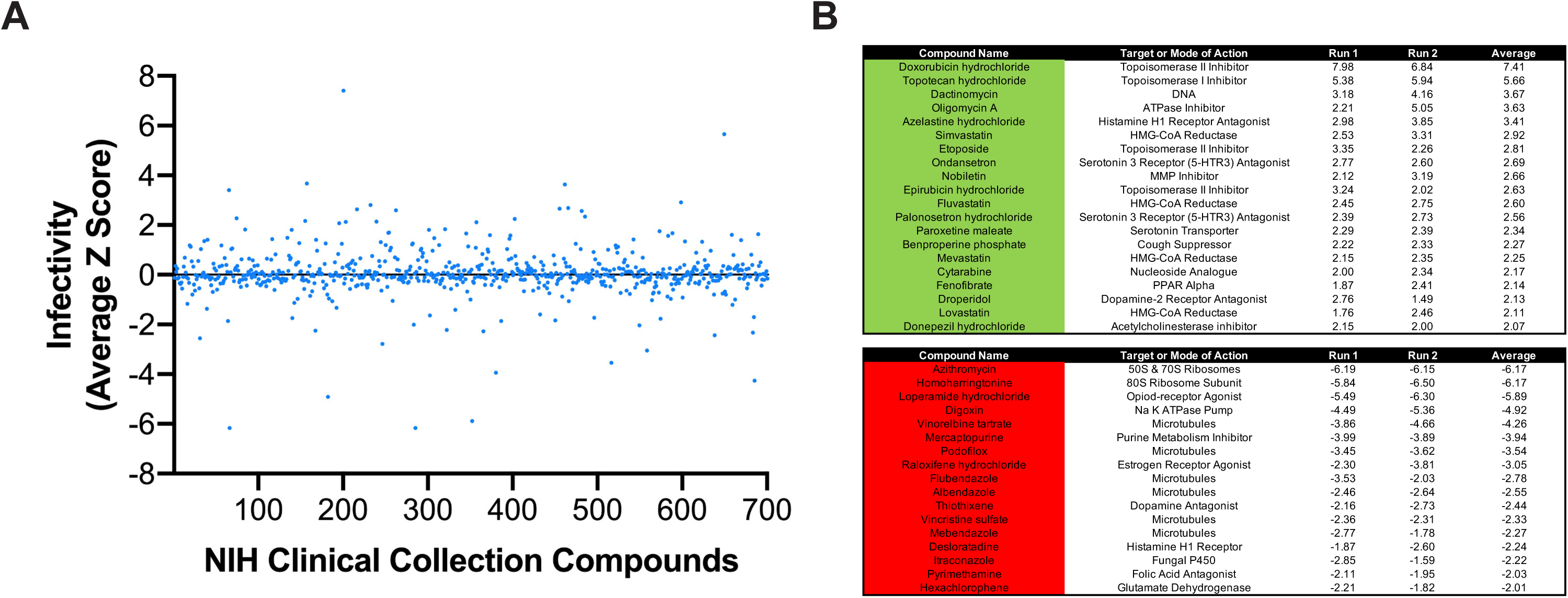
Screening of NIH Clinical Collection small molecules for reovirus infectivity. MDA-MB-231 cells were treated with vehicle (DMSO), 4 μM E64-d, or 10 μM compounds from the NIH Clinical collection for 1 h, infected with r2Reovirus at an MOI of 20 PFU/cell in the presence of DMSO, 2 μM E64-d, or 5 μM compounds from the NIH Clinical collection for 20 h. Cells were scored for infectivity by indirect immunofluorescence using reovirus-specific antisera. (A) Data are shown as infectivity from average Z-scores for compounds in the NIH Clinical Collection for duplicate experiments. (B) Compounds from the NIH Clinical Collection that increase (green, top table) or decrease (red, bottom table) infectivity by 2 Z-scores or more. Data are shown for each experimental replicate (Run).

### Topoisomerase inhibitors enhance reovirus infection of MDA-MB-231 cells

To determine if topoisomerase inhibitors affect reovirus infection of TNBC cells, MDA-MB-231 cells were treated with increasing concentrations of doxorubicin, epirubicin, and topotecan for 1 h at 37°C, adsorbed with mock or r2Reovirus at an MOI of 100 PFU/cell, and scored for infectivity by indirect immunofluorescence using reovirus-specific antiserum (FIG 7). Reovirus infectivity increased slightly when cells were treated with 0.1 µM and more significantly when treated with 1.0 µM with all three drugs. Treatment of cells with 10 μM doxorubicin or epirubicin decreased infectivity compared to 1.0 μM treatment, likely due to cellular cytotoxicity. In contrast, treatment of cells with 10 μM topotecan enhanced reovirus infectivity more than any other concentration tested. These data indicate that topoisomerase inhibitors augment reovirus infectivity in a concentration-dependent manner.

**FIG 7.**
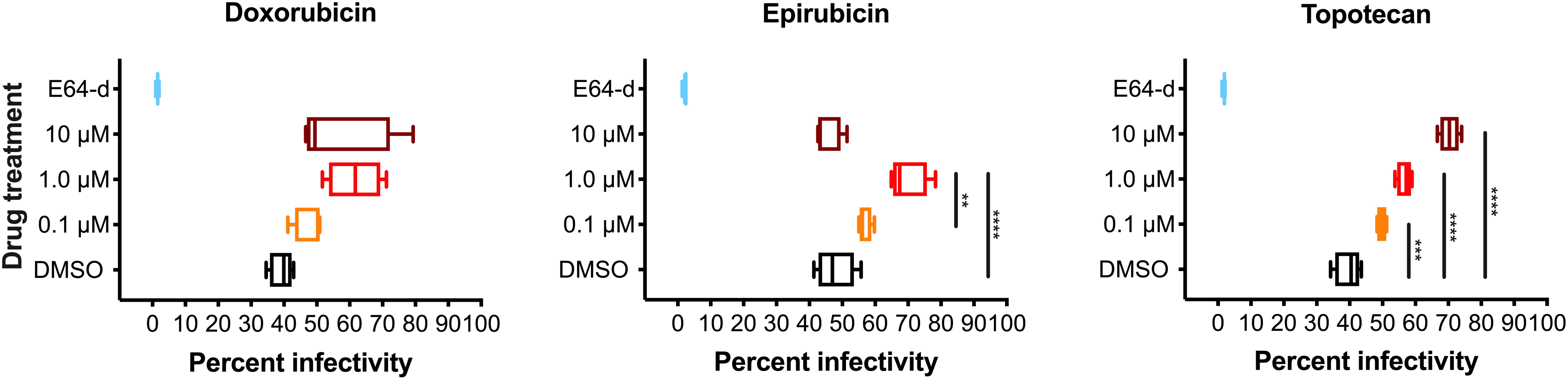
Topoisomerase inhibitors enhance reovirus infection of TNBC cells. MDA-MB-231 cells were treated for 1 h with vehicle (DMSO), 8 μM E64-d, or increasing concentrations doxorubicin, epirubicin, or topotecan and infected with r2Reovirus at an MOI of 100 PFU/cell for 20 h. Cells were assessed for infectivity by indirect immunofluorescence using reovirus-specific antisera. Data are shown as percent infectivity for quadruplicate independent experiments. **, *P* ≤ 0.01, ***, *P* < 0.001, ****, *P <* 0.0001 in comparison to DMSO by one-way ANOVA with Tukey’s multiple comparison test.

### Topoisomerase inhibitors enhance reovirus-mediated cell killing of MDA-MB-231 cells

To determine if topoisomerase inhibitors confer additive or synergistic effects on reovirus-mediated cytotoxicity, MDA-MB-231 cells were treated with vehicle (DMSO) or increasing concentrations of doxorubicin, epirubicin, or topotecan for 1 h at 37°C, adsorbed with r2Reovirus at an MOI of 200 PFU/cell, and assessed for cell viability over 3 days (FIG 8).Treatment with 0.1 μM of all three drugs did not significantly impact cell viability in the presence or absence of r2Reovirus. In the absence of virus, 1.0 μM doxorubicin and epirubicin impaired cell viability to similar levels as virus alone and moderately enhanced in the presence of reovirus. These effects can be especially observed at day 3 post infection (FIG 8B) Similar results were observed with 10 μM doxorubicin and epirubicin, except that the drugs alone had significant cytotoxic properties. In contrast, 1.0 μM topotecan had modest effects on cell viability in the absence of reovirus, but addition of reovirus had a synergistic effect on the cytotoxic effects of both topotecan and reovirus. Similar results were observed with 10 μM topotecan. Together, these data indicate that the combination of topoisomerase inhibitors with reovirus, especially topotecan, enhances the cytopathic properties of drugs and virus in a TNBC cell line.

**FIG 8.**
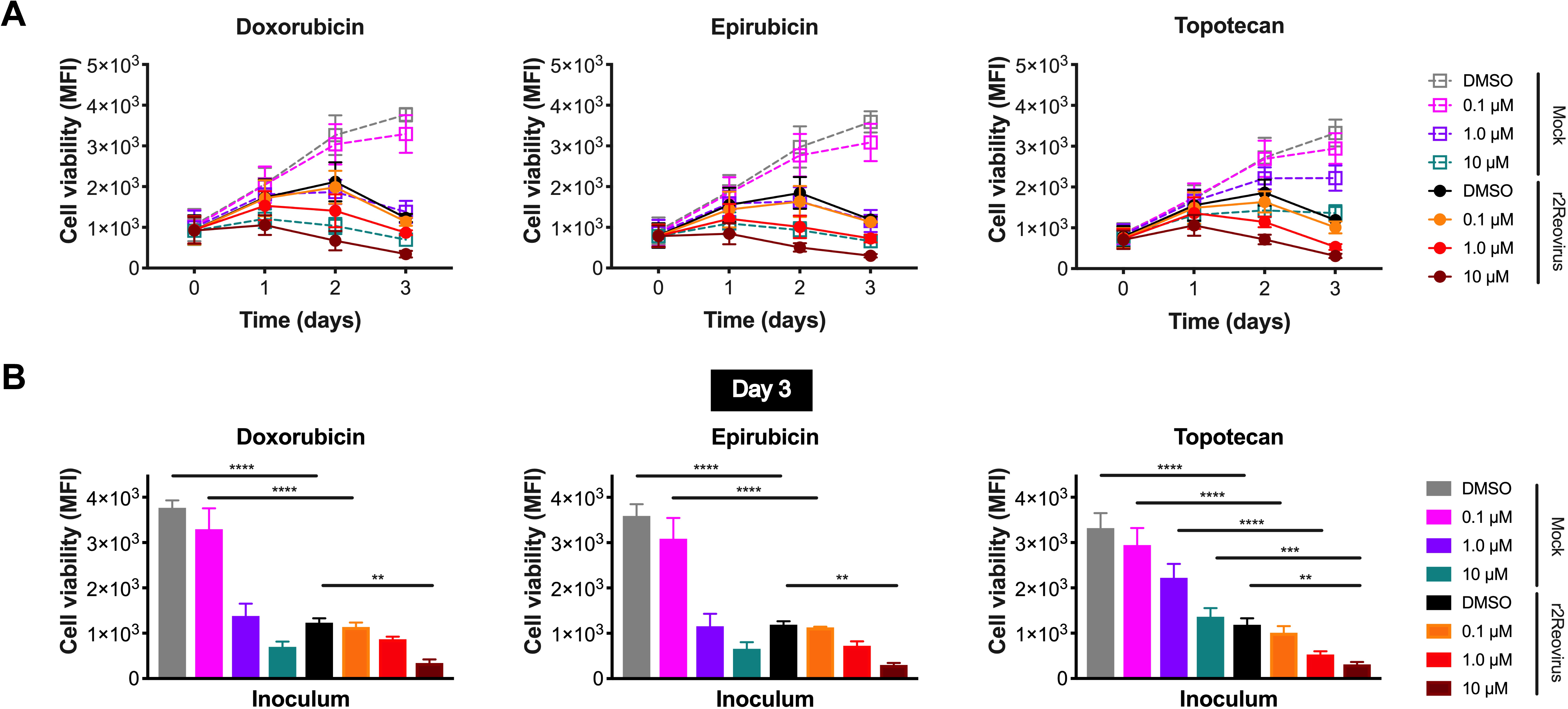
Cell viability of MDA-MB-231 cells is impaired by reovirus and topoisomerase inhibitors. (A) MDA-MB-231 cells were treated with vehicle (DMSO) or increasing concentrations of doxorubicin, epirubicin, or topotecan for 1 h, infected with r2Reovirus at an MOI of 200 PFU/cell, and assessed for cell viability at days 0-3 post infection. Data are shown as mean fluorescence intensity (MFI) for quadruplicate independent experiments. (B) Cell viability for all conditions in (A) for day 3 post infection. Error bars represent SEM. **, *P* < 0.01, ***, *P* < 0.001 by one-way ANOVA with Tukey’s multiple comparison test.

### Activation of DNA damage repair and innate immune signaling pathways following reovirus infection with topoisomerase inhibitors

Reovirus infection activates innate immune signaling that results in the production of interferon (IFN) (47, 48). Topoisomerase inhibitors, but not reovirus, induce DNA damage repair pathways and can induce innate immune signaling (49). To determine if reovirus infection of TNBC cells impacts DNA damage repair and innate immune pathways, MDA-MB-231 cells were treated with DMSO, doxorubicin, epirubicin, or topotecan for 1 h at 37°C, adsorbed with mock or r2Reovirus, whole cell lysates were collected at 0, 1, and 2 days post infection, and immunoblotted for phosphorylated and total STAT1, STAT2, STAT3, ATM, and p53 (FIG 9). Reovirus infection induced modest levels of phosphorylated STAT1 at day 1 in cells treated with DMSO and doxorubicin, but significant levels in topotecan-treated cells. Phosphorylated STAT2 was only observed in reovirus-infected topotecan-treated cells. Total levels of STAT1 and STAT2 were slightly higher in cells treated with doxorubicin, epirubicin, and topotecan compared to DMSO. STAT3 is constitutively activated in 40% of breast cancers and is associated with epithelial to mesenchymal transition (50, 51). Phosphorylated STAT3 was detected in the absence of reovirus regardless of the presence of topoisomerase inhibitors. Infection resulted in decreased levels of phosphorylated STAT3 at 1 and 2 dpi also independent of doxorubicin, epirubicin, or topotecan. These data indicate that reovirus infection of MDA-MB-231 cells promotes activation of innate immune pathways and that infection in the presence of topotecan, but not doxorubicin or epirubicin, enhances the activation of both STAT1 and STAT2. Reovirus infection also dampens the activation of STAT3 independent of topoisomerase inhibitors.

**FIG 9.**
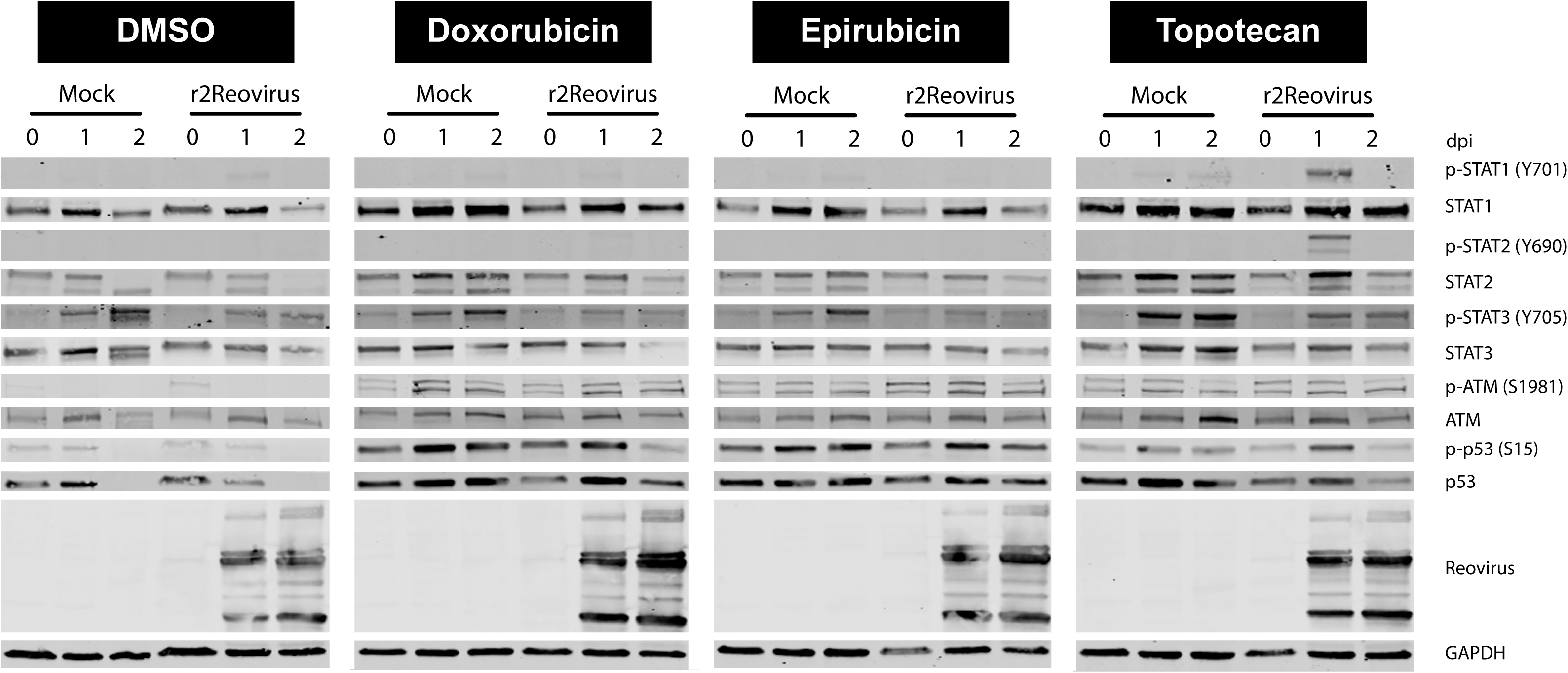
Reovirus activates STAT1 signaling and topoisomerase inhibitors activate DNA damage response pathways. MDA-MB-231 cells were treated with vehicle (DMSO) or 2 μM doxorubicin, epirubicin, or topotecan for 1 h, infected with reovirus at an MOI of 100 PFU/cell, and incubated with DMSO or 1 μM topoisomerase inhibitors for 0-3 days post infection. Whole cell lysates were resolved by SDS-PAGE and immunoblotted with antibodies specific for phosphorylated and total STAT1, STAT2, STAT3, ATM, p53 and GAPDH and reovirus. Residues recognized by phosphorylation-specific antibodies are shown in parenthesis. Results are shown for a representative immunoblot of two independent experiments.

Reovirus infection in the absence of topoisomerase inhibitors did not result in increased levels of either phosphorylated ATM or p53 compared to mock-infected cells. Treatment of cells with topoisomerase inhibitors in the absence of reovirus increased levels of phosphorylated ATM and p53 compared to DMSO-treated cells at all time points tested. The activation of ATM and p53 by topoisomerase inhibitors was not affected by the presence of reovirus. These data suggest that reovirus does not affect the activation of DNA damage signaling activated by topoisomerase inhibitors.

### Reovirus infection of TNBC cells results in increased levels of Type III interferon

To assess if the increased levels of phosphorylated STAT1 and STAT2 correlate with IFN production during reovirus infection, MDA-MB-231 cells were treated with DMSO, doxorubicin, epirubicin, or topotecan for 1 h at 37°C, infected with r2Reovirus at an MOI of 100 PFU/cell, and RNA and supernatants were collected at 0, 8, 12, 24, and 48 h post infection (FIG 10). Reovirus mRNA levels were largely unaffected by the presence or absence of topoisomerase inhibitors up to 12 h post infection and slightly increased in doxorubicin and epirubicin at 24 and 48 h post infection compared to DMSO and topotecan (FIG 10A). Despite robust infection, negligible levels of *IFNB1* mRNA were observed in the presence or absence topoisomerase inhibitors (FIG 10B). In contrast, significant levels of *IFNL1* mRNA were observed starting at 8 h post infection in infected cells and up to 48 h post infection (FIG 10C). In infected cells, *IFNL1* mRNA levels were higher in DMSO- and topotecan-treated cells at 8 and 12 h post infection than in doxorubicin- and epirubicin-treated cells, with the latter peaking at 24 h post infection. Interestingly, significant levels of *IFNL1* mRNA were observed at 24 h and 48 h in uninfected cells treated with doxorubicin and epirubicin. To determine if increasing levels of *IFNL1* mRNA result in increasing levels of protein, IFNλ levels were assessed by ELISA (FIG 10D). Secreted IFNλ was detected only in infected cells, except for low levels at 48 h in uninfected cells. IFNλ was first observed at 12 h post infection only in epirubicin-treated cells. By 24 h post infection, IFNλ was observed at similar levels in cells treated with DMSO, doxorubicin, and topotecan, but not epirubicin. At 48 h post infection, high levels of IFNλ were observed in all infected conditions, with the highest levels observed in topotecan-treated cells. These data show that topoisomerase inhibitors do not affect overall reovirus replication kinetics and that reovirus infection of MDA-MB-231 cells results in increased levels of Type III, but not Type I, IFN mRNA and protein. Although topoisomerase inhibitors had a modest effect in the induction of *IFNL1* mRNA following reovirus infection, the presence of topotecan had the largest effect on the levels of secreted IFNλ.

**FIG 10.**
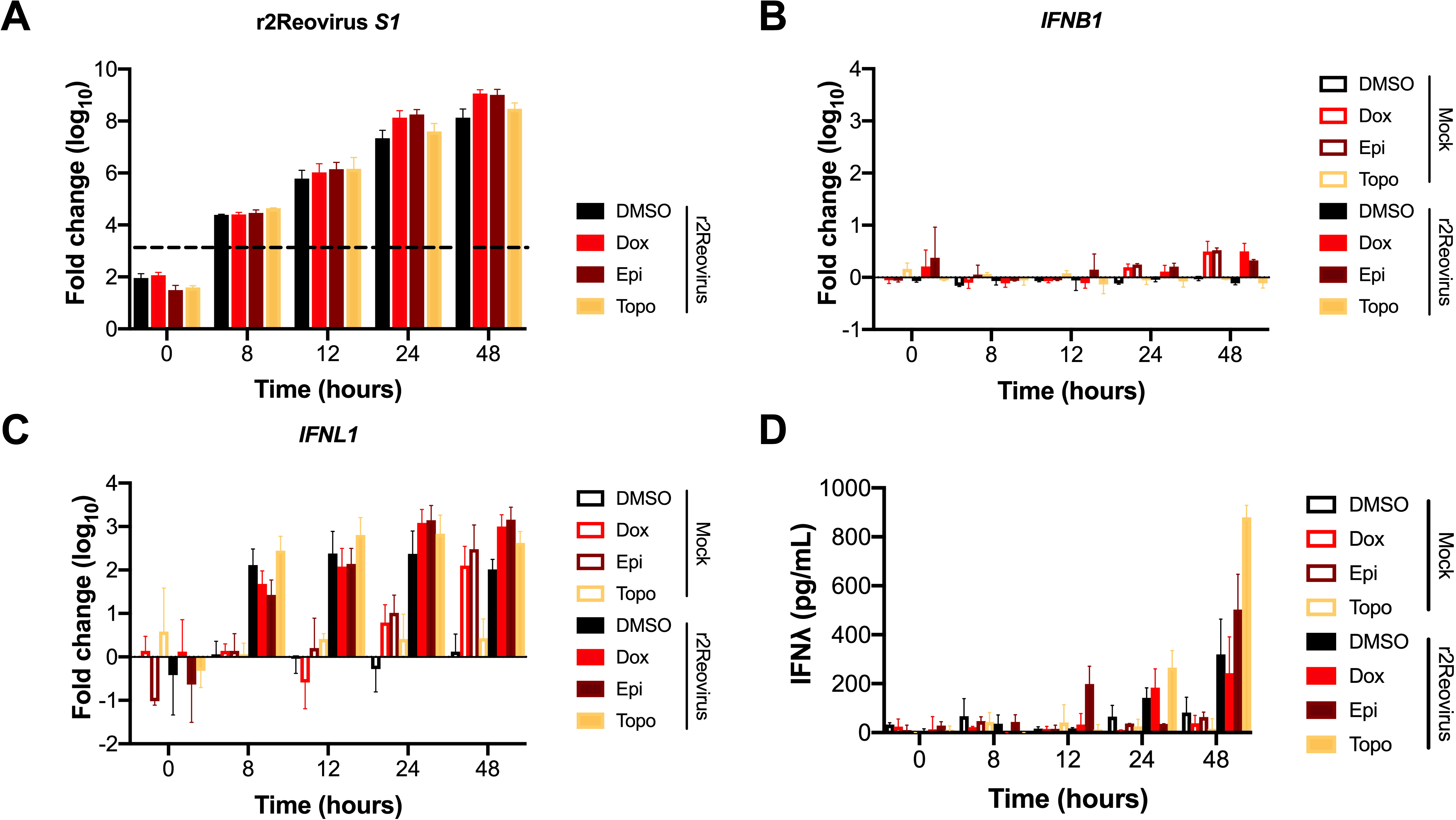
Reovirus infection of MDA-MB-231 cells induces IFNλ. MDA-MB-231 cells were treated with vehicle (DMSO) or 2 μM doxorubicin, epirubicin, or topotecan, infected with Mock or r2Reovirus at an MOI of 100 PFU/cell, and RNA (A-C) or supernatants (D) were collected at 0-48 h post infection. qPCR was performed to assess mRNA levels of (A) reovirus *S1*, (B) *IFNB1*, or (C) *IFNL1*. Data are shown as fold change normalized to housekeeping gene for duplicate independent experiments. Dashed line in (C) represents background baseline level observed in mock. (D) Levels of IFNλ in cell supernatants detected by ELISA. Data are shown as pg/ml of IFNλ for duplicate independent experiments.

## Discussion

Reovirus has an inherent preference to replicate in tumor cells, making it ideally suited for use in oncolytic therapy (15, 16). Reovirus can be delivered to patients via intratumoral and intravenous administration and can be effective in combination therapy (14). A Type 3 reovirus (T3C$) is currently in Phase I-II clinical trials against a variety of cancers in combination with several drugs (clinicaltrials.gov: NCT01622543, NCT01656538). In this study, we generated novel reassortant reoviruses with enhanced replicative properties in TNBC cells by coinfection of a TNBC cell line with prototype strains T1L, T2J, and T3D and serial passage. Reassortant reoviruses attach to cells with similar efficiency as T1L, whereas Type 3 reoviruses attach with enhanced efficacy. T1L uses GM2 glycans to attach to cells whereas T3D interacts with α2,3-linked sialic acid (39, 52). High expression of α2,3-sialic acid in breast cancer is associated with greater metastatic potential (53), suggesting the slight enhancement in attachment observed with Type 3 reoviruses could be due to high levels of α2,3-sialic acid present on the surface of MDA-MB-231 cells.

Reassortant viruses did not have mutations in σ1 and the most predominant viruses following serial passaging all had a Type 1 σ1. These data suggest that carbohydrate binding did not drive selection of the reassortant viruses. JAM-A is expressed in normal mammary epithelial cells and high JAM-A expression in breast cancer patients correlates with worse survival and increased recurrence (54, 55). MDA-MB-231 cells express JAM-A (55), although relatively low JAM-A levels may be responsible for the lower infectivity observed by all reoviruses tested in comparison to infection in L929 cells. These data suggest that receptor engagement is not responsible for the enhanced infectivity observed with the reassortant viruses.

During cell entry, reovirus traverses to endosomes where cathepsin proteases cleave outer capsid protein σ3, forming an infectious subvirion particle (ISVP) (41, 56). Both reassortants have a nonsynonymous mutation in the σ3-encoding S4 gene segment that results in a V49I substitution. This mutation has not been identified to impact reovirus disassembly kinetics, but it is possible it could expedite viral cell entry kinetics. However, reassortant viruses were equally sensitive to E64-d treatment as parental viruses. Although reassortant viruses infected MDA-MB-231 cells more efficiently than T1L, T3D, and T3C$, replication kinetics of the reassortant viruses were similar except for T3D, which had slower replication kinetics. These data indicate that Type 1 reoviruses replicate with enhanced kinetics compared to T3D, but that genetic differences between T3D and T3C$ are sufficient to allow T3C$ to replicate as efficiently as Type 1 viruses. These data also suggest that the enhanced cytotoxic properties of the reassortant viruses over parental viruses are not due to enhanced replication kinetics in MDA-MB-231 cells.

The reovirus L3, S2, and S3 gene segments have distinct roles in reovirus replication. The L3-encoded λ1 protein is a major inner-capsid protein that has phosphohydrolase activity and participates in viral transcription (57, 58). The S2-encoded σ2 protein is essential for the assembly of viral cores (59). The S3-encoded nonstructural protein σNS is required for viral factory formation (60). The similarity in replication efficiency observed between T1L and the reassortant viruses suggests the A160T mutation in L3 and I250V mutation in S3 (found in both reassortants) and the V426I in S2 and P161T in S3 (in r1Reovirus only) do not impact overall replication efficiency. However, it is possible that point mutations in these gene segments in the reassortant viruses impact the activity of the viral proteins that result in enhanced infectivity or cytotoxicity in the context of TNBC cells.

Of all the viruses tested in MDA-MB-231 cells, r1Reovirus and r2Reovirus impaired cell viability with the fastest kinetics, and only T3D was severely deficient in killing these cells. The poor induction of cell death by T3D may be related to its dampened replication in these cells.

Differences in the induction of apoptosis by reovirus strains segregate with the M2 and S1 gene segments (33). Apoptosis is activated by fragments of the M2-encoded μ1 protein generated during reovirus cell entry (28, 32, 33, 61, 62). The μ1 protein impacts reovirus infectivity by enhancing reovirus attachment to cells (63). S1 is genetically linked to reovirus induction of apoptosis through the activities of both σ1 and σ1s, although it is unclear if the effects of σ1s on the induction of cell death are independent of its ability to regulate viral protein synthesis and induce cell cycle arrest (64, 65). We did not observe significant levels of cell cycle arrest in MDA-MB-231 cells infected with reassortant reoviruses (data not shown). It is unclear if the enhanced cytopathic properties of reassortant viruses in the context of TNBC cells maps to the T3D M2 gene segment, the various nonsynonymous changes, or a combination of both.

Screening small molecules from the NIH Clinical Collection identified 20 molecules that increase infectivity and 17 molecules that decreased infectivity in MDA-MB-231 cells. Six microtubule-inhibiting drugs, digoxin, and two serotonin antagonists affected reovirus infectivity, corroborating the role of microtubules, the sodium-potassium ATPase pump, and serotonin receptors in reovirus infection (43–45). Of the 17 molecules that enhanced infectivity, 4 are topoisomerase I (topotecan) or II (doxorubicin, epirubicin, and etoposide) inhibitors. Treatment of cells with topoisomerase inhibitors resulted in increased infectivity, with no effect on virus attachment (data not shown), and no significant increase in viral RNA levels at 8 and 12 h post infection and slight increases in viral RNA at 24 and 48 h post infection. Topoisomerase inhibitors promote DNA double-strand breaks leading to cell death (66–69). Reovirus infection does not induce DNA double-strand breaks and promotes cell death through the induction of extrinsic and intrinsic apoptosis or necroptosis (23, 24, 28, 33, 70, 71). It is possible that topoisomerase inhibitors positively affect uptake of viral particles during cell entry that results in enhanced infectivity and that doxorubicin and epirubicin further impact a step late in the viral life cycle that results in enhanced transcription of viral RNA. It is also possible that the additive cytotoxicity observed in MDA-MB-231 cells when both reovirus and topoisomerase inhibitors are present is through the activation of complementary cell death pathways.

Reovirus infection does not impair the DNA double strand break response activated by treatment with topoisomerase inhibitors. Late during infection in the presence of topoisomerase inhibitors, levels of phosphorylated and total p53 were lower than in uninfected cells. It remains to be determined if the effects of reovirus infection on p53 are at the transcriptional, translational, or post-translational level. Reovirus infection can induce higher levels of activated MDM2, which leads to p53 degradation (72). In the context of reovirus infection, it is possible that topoisomerase inhibitors promote p53 stabilization through impairing the activation of MDM2 by the virus. It is also possible the effects on total p53 at late times post infection are due to viral-dependent host translational shutoff. In support of this, total levels of STAT1, STAT2, STAT3, and ATM were also lower at late times of infection.

Reovirus infection of MDA-MB-231 cells resulted in robust expression of Type III, but not Type I, IFN mRNA and protein. Infection in the presence of topoisomerase inhibitors did not significantly affect levels of *IFNL1* mRNA. Interestingly, doxorubicin and epirubicin treatment in the absence of infection results in the induction of *IFNL1* mRNA starting at 24 h reaching similar levels to those detected in virus-infected cells by 48 h. Induction of DNA double strand breaks by topoisomerase inhibitors can result in p53-dependent regulation of Type I IFN through a STING-dependent but cGAS-independent pathway (49). MDA-MB-231 cells express STING (data not shown), suggesting that topoisomerase inhibitors could be inducing transcription of Type III IFN downstream of the induction of the DNA damage response through a similar mechanism. However, topoisomerase inhibitors did not induce Type I IFN transcription in the presence or absence of reovirus.

Levels of IFNλ were first observed at 12-24 h post infection in the presence or absence of topoisomerase inhibitors, with the highest levels of IFNλ detected at 48 h post infection in the presence of topotecan. IFN-λ1, IFN-λ2, and IFN-λ3 are expressed in breast cancer cells, although their role in mediating innate immunity in these cells is not well characterized (73). Type I and Type III IFN are transcriptionally regulated by the transcription factor IRF3 (74, 75). Reovirus can antagonize IFN production by sequestering IRF3 to viral inclusions (76) and infection of gut epithelial cells *in vitro* and *in vivo* results in upregulated levels of IFNλ mRNA (76–78). It is possible that in MDA-MB-231 cells reovirus is unable to sequester IRF3 to viral inclusions, resulting in robust production of Type III IFN. Reovirus infection of TNBC cells resulted in high levels of secreted IFNλ, with over 200 pg/ml detected at 48 h post infection in the presence or absence of topoisomerase inhibitors. Levels of IFNλ in the presence of topotecan at 48 h post infection reached over 800 pg/ml, levels that are higher than that observed in dendritic cells that have been exposed to a RIG-I agonist (79). It is unclear why topotecan, but not doxorubicin or epirubicin result in significantly higher IFNλ levels, especially considering that *IFNL1* mRNA levels were not different in cells infected in the presence of the different topoisomerase inhibitors. MDA-MB-231 cells can express low basal levels and are responsive to Type I IFNs (80–82). The large levels of Type III IFN detected in MDA-MB-231 cells, and lack of Type I IFN, indicates that STAT activation observed in these cells is likely in response to the interaction of IFNλ with its receptors.

Despite the robust induction of Type III IFN in response to infection, robust levels of activated STAT1 and STAT2 were only detected in the presence of topotecan. Low levels of activated STAT1 were observed in infected cells in the absence of topoisomerase inhibitors, but no STAT activation was observed in the presence of doxorubicin or epirubicin. It is possible that the low levels of activated STAT1 and STAT2 in infected MDA-MB-231 cells are a result of impaired sensing of IFNλ due to low level expression of the IFNλ receptor. It is also possible that treatment of cells with topotecan may sensitize cells to IFNλ through the upregulation of the IFNλ receptor. Surprisingly, despite high levels of activated STAT1 and STAT2 following reovirus infection of topotecan-treated cells, reovirus infectivity and replication remained unimpaired.

In this study, we generated reoviruses with unique infective and cytotoxic properties by forward genetics following coinfection with three different serotype reoviruses. The novel genetic composition of the reassortant viruses could inform future studies on viral factors that promote infection and killing of cells by reovirus. Through high-throughput screening we identified topoisomerase inhibitors as a class of drug that enhances infection and the cytotoxic properties of reovirus in the context of TNBC. We also show that infection of a breast cancer cell line leads to the robust production of Type III, but not Type I, IFN. This study presents evidence for the pairing of reassortant reoviruses generated by forward genetics with topoisomerase inhibitors identified by high-throughput screening as a promising therapeutic against TNBC.

## Materials and Methods

### Cells, viruses, and antibodies

MDA-MB-231 cells (gift from Jennifer Pietenpol, Vanderbilt University) and MDA-MB-436 cells (ATCC HTB-130) were grown in Dulbecco’s Modified Eagle’s Medium (DMEM) supplemented with 10% fetal bovine serum (FBS) (Life Technologies), 100 U per ml penicillin and streptomycin (Life Technologies). Spinner-adapted L929 cells (Terry Dermody, University of Pittsburgh) were grown in Joklik’s modified MEM with 5% FBS, 2 mM L-glutamine (Life Technologies), penicillin and streptomycin, and 0.25 mg per ml amphotericin B (Life Technologies).

Reovirus strains Type 1 Lang (T1L) and Type 3 Dearing (T3D) working stocks were prepared following rescue with reovirus cDNAs in BHK-T7 cells (gift from Terry Dermody, University of Pittsburgh), followed by plaque purification, and passage in L929 cells (83). Reovirus type 2 Jones (T2J) is a laboratory strain and Type 3 Cashdollar (T3C$) is a distinct Type 3 reovirus (84). Purified virions were prepared using second-passage L929 cell lysate stocks. Virus was purified from infected cell lysates by Vertrel XF (TMC Industries Inc.) extraction and CsCl gradient centrifugation as described (85). The band corresponding to the density of reovirus particles (1.36 g/cm^3^) was collected and dialyzed exhaustively against virion storage buffer (150 mM NaCl, 15 mM MgCl2, 10 mM Tris-HCl [pH 7.4]). Reovirus particle concentration was determined from the equivalence of 1 unit of optical density at 260 nm to 2.1×10^12^ particles (86). Viral titers were determined by plaque assay using L929 cells (87). Reovirus virions were labeled with succinimidyl ester Alexa Fluor 488 (A488) (Life Technologies) as described (43, 88).

Reovirus polyclonal rabbit antiserum raised against reovirus strains T1L and T3D was purified as described (89) and cross-adsorbed for MDA-MB-231 cells. Secondary IRDye 680 and 800 antibodies (LI-COR Biosciences) and goat anti-rabbit Alexa Fluor 488 (A488) (Life Technologies).

### Serial passage of T1L, T2J, and T3D in MDA-MB-231 cells

MDA-MB-231 cells were adsorbed with T1L, T2J, and T3D at a multiplicity of infection (MOI) of 1 PFU/cell for 1 h at room temperature and incubated for 48 h at 37°C in MDA-MB-231 cell media. Cells were freeze-thawed three times, fresh MDA-MB-231 cells were infected with 500 μl of freeze-thawed cell supernatant, and incubated for 48 h at 37°C. Serial passage was repeated 20 times and individual viral titers were obtained by plaque isolation following plaque assay in L929 cells.

### Electrophoretic mobility of reovirus

5×10^10^ particles of purified reovirus or freeze-thawed supernatants containing reovirus mixed with 2X SDS-Sample Buffer (20% Glycerol, 100 mM Tris-HCl [pH 6.8], 0.4% SDS, and 3 mg Bromophenol Blue) were separated by SDS-PAGE using 4-20% gradient polyacrylamide gels (Bio-Rad Laboratories) at 10 mAmps for 16 h. The gel was stained with 5 μg/ml ethidium bromide for 20 min and imaged using a Chemidoc XRS+ (Bio-Rad).

### Next Generation Sequencing of Reovirus

RNA from viral preparations of T1L, T2J, T3D, r1Reovirus, and r2Reovirus were obtained using an RNeasy RNA purification kit (Qiagen). Ten nanograms of viral RNA was used as input for cDNA synthesis using the Clontech SMARTer Stranded Total RNA-Seq Kit v2 (Pico Input, Mammalian) according to the manufacturer’s instructions. Libraries were validated by capillary electrophoresis on an Agilent 4200 TapeStation, pooled, and sequenced on an Illumina HiSeq3000 with 100bp paired end reads averaging 13 million reads/sample, yielding an average depth of coverage > 1000 reads. Reads were trimmed of adapter sequence using Trimmomatic (version 0.36, http://www.usadellab.org/cms/?page=trimmomatic) using the TruSeq3-PE-2 paired end adapter reference. Trimmed reads from each sample were aligned to all of the parental strain reference sequences using the Burrows-Wheeler Aligner (BWA version 0.7.10-r789, http://bio-bwa.sourceforge.net/). Deduplication was performed with Picard tools (version 1.74(1243), https://broadinstitute.github.io/picard/), and variation was called, again for each sample against all the parental strain references, using the GATK pipeline’s (version 3.4, https://software.broadinstitute.org/gatk/) HaplotypeCaller with ploidy set to 1 and other default parameters. The resultant Variant Call Files (.vcf) were examined for sample similarity/variation from the parental reference strains. The read files for this study have been deposited with the NCBI Sequence Read Archive (SRA) and are available via the accession (In process).

### Flow cytometric analysis of cell-surface reovirus

MDA-MB-231 cells were adsorbed with 5×10^3^-5×10^4^ particles per cell of A633-labeled virus for 1 h at room temperature. Cells were washed with PBS, detached with Cellstripper (Cellgro) for 10 min at 37°C, quenched and washed with PBS containing 2% FBS. Cells were fixed in 1% EM-grade paraformaldehyde (Electron Microscopy Sciences). Mean fluorescence intensity (MFI) was assessed using a CytoFLEX flow cytometer (Beckman Coulter) and quantified using FlowJo software.

### Reovirus infectivity assay

Reovirus infectivity was assessed by indirect immunofluorescence (90). MDA-MB-231 and L929 cells were adsorbed with reovirus at a range of MOIs for 1 h at room temperature, washed with PBS, and incubated in media for 16-24 h at 37°C. To assess the effects of topoisomerase inhibitors on reovirus infectivity, cells were pretreated with topoisomerase inhibitors or E64-d for 1 h at 37°C, reovirus was added to cells, and incubated for 18-24 h at 37°C. Cells were fixed with ice-cold methanol and stored at -20°C for at least 30 min. Methanol was removed, cells were washed twice with PBS, and blocked with PBS containing 1% BSA for 15 min at room temperature. Cells were stained with reovirus-specific polyclonal antiserum (1:2000) for 1 h at room temperature, washed twice with PBS, stained with goat anti-rabbit Alexa 488 (1:1000) for 1 h at room temperature, counterstained with 0.5 ng/ml DAPI for 5 min at room temperature, and washed twice with PBS. Immunofluorescence was detected using a Lionheart FX Automated Microscope (Biotek) with a 4x-PLFL phase objective (NA 0.13), and percent infectivity was determined (reovirus positive cells/DAPI positive cells) using Gen5 software (Biotek).

### Reovirus replication assay

MDA-MB-231 cells were adsorbed with reovirus at a MOI of 10 PFU/cell for 1 h at room temperature, washed with PBS, and incubated for 0-3 days in MDA-MB-231 media at 37°C. Cells were freeze-thawed three times and viral titers were determined by plaque assay using L929 cells. Viral yields were calculated by dividing viral titers by the viral titer from day 0.

### Cell viability assay

Cell viability was assessed by measuring metabolic activity using Presto Blue reagent (Invitrogen). L929, MDA-MB-231, and MDA-MB-436 cells were adsorbed with reovirus at a range of MOIs for 1 h at room temperature or treated with 1 μM staurosporine, washed with PBS, and incubated for 0-7 days at 37°C. To determine the effects of topoisomerase inhibitors on cell viability, cells were pretreated with increasing concentrations of topoisomerase inhibitors for 1 h at 37°C, reovirus was added to cells, and incubated in the presence of the inhibitors for 0-3 days. Presto Blue was added at each time point for 30 min at 37°C and fluorescence (540 nm excitation/590 nm emission) was measured with a Synergy HT plate reader (Biotek).

### Screening of NIH Clinical Collection Small Molecule Inhibitors

The NIH Clinical Collection was obtained from the NIH Roadmap Molecular Libraries Screening Centers Network. MDA-MB-231 cells were treated with DMSO, 4 μM E64-d, or 10 μM of compounds from the NIH Clinical Collection for 1 h at 37°C. Media (mock) or reovirus was added to cells at an MOI of 20 PFU/cell, and incubated for 20 h at 37°C. Cells were fixed and scored for infectivity by indirect immunofluorescence as described previously. Z scores for each well were calculated using the following formula: Z score = (a-b)/c, where a is the percent infectivity (infected cells/number of cells), b is the median percent infectivity for each plate, and c is the standard deviation of percent infectivity for each plate. Z scores of -2 > x < 2.0 were considered significant. Data for all compounds in screen are provided in Table S2.

### Immunoblotting for DNA damage response and innate immune molecules

MDA-MB-231 cells were treated with DMSO or 2 μM topoisomerase inhibitors for 1 h at 37°C, infected with mock or reovirus at an MOI of 100 PFU/cell, and incubated for 0-2 days at 37°C. Whole cell lysates were prepared using RIPA buffer (20 mM Tris-HCl [pH 7.5], 150 mM NaCl, 1 mM EDTA, 1% NP-40, 0.1% sodium dodecyl sulfate, 0.1% sodium deoxycholate) and fresh Protease Inhibitor Cocktail (P8340, Sigma-Aldrich), Phosphatase Inhibitor Cocktail 2 (P5726, Sigma-Aldrich), 1 mM sodium vanadate, and 1 mM phenylmethylsulfonyl fluoride (PMSF) and protein concentration was determined using the DC protein assay (Bio-Rad). Whole cell lysates were resolved by SDS-PAGE in 4-20% gradient Mini-PROTEAN TGX gels (Bio-Rad) and transferred to 0.2 μm pore size nitrocellulose membranes (Bio-Rad). Membranes were incubated for 1 h in blocking buffer (Tris-buffered saline [TBS] with 5% powdered milk), incubated with primary antibodies specific for phospho-STAT1 (Y701, clone D4A7 #7649), - STAT2 (Y690, clone D3P2P, #88410), -STAT3 (Y705, clone D3A7, #9145), -ATM(S1981, clone 10H11.E12, #4526), -p53(S15, #9284), total STAT1 (clone D3A7, #9145), STAT2 (clone D9J7L, #72604), STAT3 (clone 124H6, #9139), ATM (clone D2E2, #2873), p53 (clone 1C12, #2524), and GAPDH (clone GA1R, MA5-15738), and reovirus polyclonal antiserum overnight at 4°C. Antibodies are from Cell Signaling Technology except for GAPDH, ThermoFisher. Membranes were washed with TBS-T (TBS with 0.1% Tween 20) and incubated with secondary antibodies conjugated to IRDye 680 or IRDye 800. Membranes were imaged using a LiCor Odyssey CLx and processed in ImageStudio (LI-COR Biosciences).

### qPCR assessment of Type 1 and 3 interferon transcript levels

MDA-MB-231 cells were treated with DMSO or 2 μM topoisomerase inhibitors for 1 h at 37°C, infected with mock or r2Reovirus at an MOI of 100 PFU/cell, and incubated for 0, 8, 12, 24, and 48 h. RNA was isolated using a QIAGEN RNeasy kit with on-column DNase digestion. cDNAs were generated using 500 ng of RNA and random primers with the High-Capacity cDNA Reverse Transcription Kit (ThermoFisher) in a SimpliAmp Thermal Cycler (ThermoFisher). cDNA was diluted 1:5 in nuclease-free water and qPCR reactions were performed in MicroAmp Fast Optical 96-Well Reaction Plates (Applied Biosystems) using PrimeTime qPCR assays (IDT) for *IFNB1*, *IFNL1*, *HPRT1*, and a custom assay for the reovirus S1 gene segment (<u>Probe</u>: 5’-/56-FAM/TCAATGCTG/ZEN/TCGAACCACGAGTTGA/3IABkFQ/-3’, <u>Primer 1</u>: 5’-CGAGTCAGGTCACGCAATTA-3’; <u>Primer 2</u>: 5’-GGATGTTCGTCCAGTGAGATTAG-3’) using a 7500 Fast Real-Time PCR System (Applied Biosystems) and accompanying software to analyze qPCR data.

### IFNλ ELISA

MDA-MB-231 cells were treated with DMSO or 2 μM topoisomerase inhibitors for 1 h at 37°C, infected with mock or r2Reovirus at an MOI of 100 PFU/cell, and incubated for 0, 8, 12, 24, and 48 h. Cell supernatants were collected and levels of IFNλ were determined with the IFN-lambda 1/3 DuoSet ELISA kit (R&D Systems). Plates were read on a Synergy HT plate reader (Biotek) using 450 nm for sample detection and 540 nm for wavelength correction.

### Statistical analysis

Mean values for quadruplicate experiments were compared using one or two-way analysis of variance (ANOVA) with Tukey’s multiple-comparison test (Graph Pad Prism). *P* values of < 0.05 were considered statistically significant.

## Acknowledgements

We would like to thank Aspen Hirsch for review of the manuscript and suggestions. This work was supported by funding from the Children’s Healthcare of Atlanta and the Pediatric Research Institute and Winship Comprehensive Cancer Institute #IRG-14-188-01 from the American Cancer Society (B.A.M.). Flow cytometry experiments were performed in the Emory Pediatrics Flow Cytometry Core (UL1TR002378). Imaging was performed at the Emory Integrated Cellular Imaging Core (2P30CA138292-04 and the Emory Pediatrics Institute). The Yerkes NHP Genomics Core is supported in part by NIH P51 OD011132. The funders had no role in the study design, data collection and analysis, decision to publish, or preparation of manuscript.

